# Age-related Changes in Memory for Object and Position-In-Context

**DOI:** 10.1101/2020.11.22.393355

**Authors:** Tammy Tran, Kaitlyn E. Tobin, Sophia H. Block, Vyash Puliyadi, Michela Gallagher, Arnold Bakker

## Abstract

There has been considerable focus on investigating age-related memory changes in cognitively healthy older adults, in the absence of neurodegenerative disorders. Previous studies have reported age-related domain-specific changes in older adults, showing increased difficulty encoding and processing object information but minimal to no impairment in processing spatial information compared to younger adults.

However, few of these studies have examined age-related changes in the encoding of concurrently presented object and spatial stimuli, specifically the integration of both spatial and non-spatial (object) information. To more closely resemble real-life memory encoding and the integration of both spatial and non-spatial information, the current study developed a new experimental paradigm with novel environments that allowed for the placement of different objects in different positions within the environment. The current findings show that older adults have decreased performance in recognizing changes of the object position within the spatial context but no significant differences in recognizing changes in the identity of the object within the spatial context compared to younger adults. These findings suggest there may be potential age-related differences in the mechanisms underlying the representations of complex environments and furthermore, the integration of spatial and non-spatial information may be differentially processed relative to independent and isolated representations of object and spatial information.

## Introduction

Advancing age is associated with changes in a number of cognitive domains (Craik & Bialystok, 2006; Erickson & Barnes, 2003; Salthouse, 2004) with particular focus and attention examining age-related changes in episodic memory function (Craik & Bialystok, 2006; Grady & Craik, 2000). In addition to a general decline in long-term retention, older adults show a reduced ability to differentiate between highly similar object representations (Berron et al., 2018; Reagh et al., 2016; Stark et al., 2013, 2015; Yassa et al., 2011) and object features (Yeung et al., 2017) compared to young adults. In contrast, spatial representations appear to be relatively spared (Fidalgo et al., 2016; Stark & Stark, 2017) with older adults showing performance similar to young adults recognizing subtle changes in a spatial environment (Berron et al., 2018) and changes in the location of an object when presented on a blank screen (Reagh et al., 2016, 2018).

The dissociation and integration of object and spatial information has been a key question in memory research and has been hypothesized to arise from two parallel information processing streams in the medial temporal lobe (Davachi, 2006; Eichenbaum, 1999; Eichenbaum et al., 1999; Knierim et al., 2013; Ranganath & Ritchey, 2012). One pathway, commonly referred to as the “what” pathway involves the perirhinal cortex and the lateral entorhinal cortex, and is thought to predominately process information about objects, items and events, while the “where” pathway with the parahippocampal cortex and the medial entorhinal cortex is thought to process contextual and spatial information. Information from both pathways is projected to the hippocampus, which is then thought to integrate the spatial and non-spatial information into a cohesive “memory space” through a mechanism that is common to both object or episodic and spatial information (Eichenbaum et al., 1999; 2014).

However, emerging evidence suggests these processing of object and spatial information may be more integrated than previously thought. In rodent studies, the lateral entorhinal cortex has been reported to be involved in the encoding of features from both the object and the environment (Deshmukh et al., 2012; Deshmukh & Knierim, 2011; Knierim et al., 2013; Yoganarasimha et al., 2011). Rodent studies using single cell recordings in the lateral entorhinal cortex show that neurons in this region encode object-related information as well as spatial information about the object (e.g., position in relationship to the environment). These studies also show cells in the lateral entorhinal cortex that track the position of an object in the environment and do not fire when that object is no longer present (Deshmukh & Knierim, 2011). A different subset of cells (‘object trace cells’) have been reported to fire in previously experienced positions of an object within an environment (Deshmukh & Knierim, 2011; Tsao et al., 2013). Lateral entorhinal cortex lesioned rodents show no impairment in performing an object-recognition task, but are impaired at recognizing spatial changes and object changes within a set of objects in an environment including position changes of the objects (Van Cauter et al., 2013) and previously learned object-place and object-context associations (Chao et al., 2016; Wilson, Langston, et al., 2013). Together, these findings suggest that, beyond encoding information about objects, the lateral entorhinal cortex encodes contextual information and may be binding non-spatial and spatial information, specifically encoding information about objects and certain spatial properties, including information about the object’s position within environment.

Few studies have examined the role of the lateral entorhinal cortex in encoding object identity, object position or changes to the spatial context in humans. Reagh and colleagues (2014) report that subtle perceptual differences between similar objects (e.g., two slightly different apples) elicits activity observed with functional magnetic resonance imaging (fMRI) in both the lateral entorhinal cortex and perirhinal cortex, while changes to an object’s position on a blank screen elicited activity in both the parahippocampal cortex and medial entorhinal cortex. Subsequent studies in older adults show impaired performance recalling the identity of object (Berron et al., 2018; Reagh et al., 2018; Reagh & Yassa, 2014; Stark & Stark, 2017; Yeung et al., 2017) but similar performance recalling position of the object on a screen compared to young adults (Reagh et al., 2016, 2018). However, these studies examined memory for object identity and object position on a blank screen, devoid of any spatial or contextual information. Given the findings from rodent studies, it appears that object identity and object position information are represented in relationship to the spatial environment in which they occur.

To examine the integration of non-spatial and spatial information, novel stimuli were developed to mimic real-life environments with different positions where objects could occur within the environment, more closely resembling animal studies in which rodents experience objects within an environment. These stimuli were employed in a novel object-in-context task to assess memory for object identity and object position in context and examine age-related changes in performance on this task in cognitively normal older adults compared to young adults.

## Methods

### Experiment 1: Scene Ratings

#### Participants

A total of 181 Johns Hopkins undergraduate students (108 females; 73 males) participated in the experiment in exchange for course credit. Data from seventeen participants were removed from analysis due to inability or failure to complete the experiment while data from three participants were removed for a nonresponse rate above 20% and data from four participants were removed for below chance accuracy. Complete data from 157 (90 females; 67 males, aged 18 - 22 years old) participants were included in the final analysis for the scene rating experiment.

#### Materials and Procedures

A total of 549 indoor scenes were created using the Sims 4 computer game (EA Games). All scenes were designed to have the same perspective, spatial dimensions and outdoor scenery (Figure 1A). Scenes featured different types of windows, wallpaper and flooring and included sparse furniture, including chairs, tables, sofas and beds. Scenes were classified into 5 general categories: living room, dining room, kitchen, bedroom and office rooms to allow for the placement of categorically relevant furniture (Figure 1B). Critically, within each scene, 2-5 different positions were defined that an object could logically occupy (Figure 1C). These general positions for object placement were consistent across scenes.

**Figure 1.**
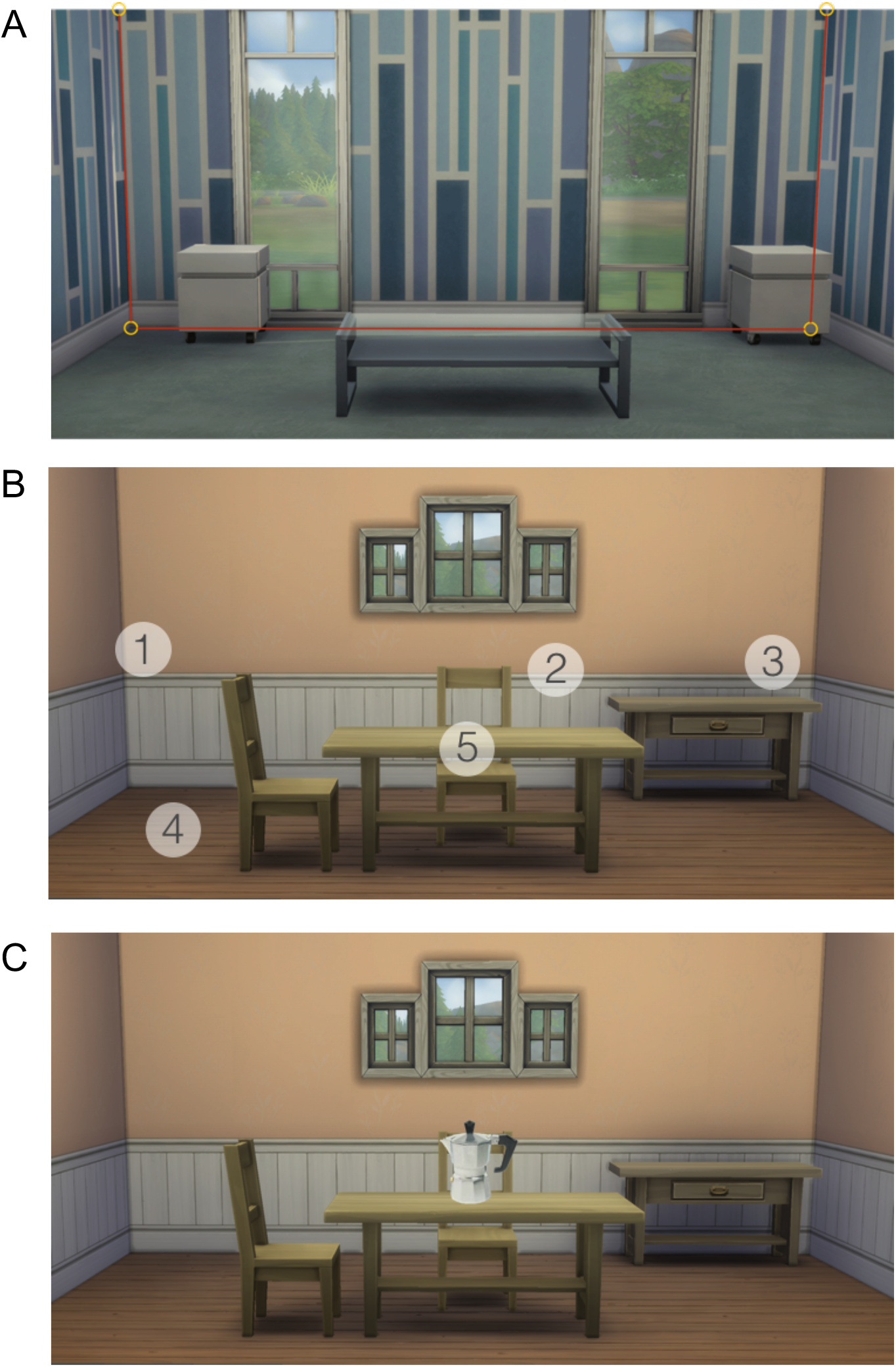
Task stimuli. A) Scenes were designed to have identical dimensions and a similar perspective. B) Scenes were designed to have 2-5 different positions where an object could be reasonably placed. Across all scenes, the same general positions were available for object placement. C) Example of a scene with an object as seen by the participant.

To examine the mnemonic attributes of the indoor scenes, participants were asked to judge whether each scene was either ‘new’ or ‘old.’ A scene was correctly judged ‘new’ if it was seen for the first time in the context of the task, and ‘old’ if the exact scene was repeated once or twice (Figure 2A). A total of 182 scenes were presented once, 245 scenes were presented twice, and 122 scenes were presented three times in random order, with participants completing a total of 549 scenes.

**Figure 2.**
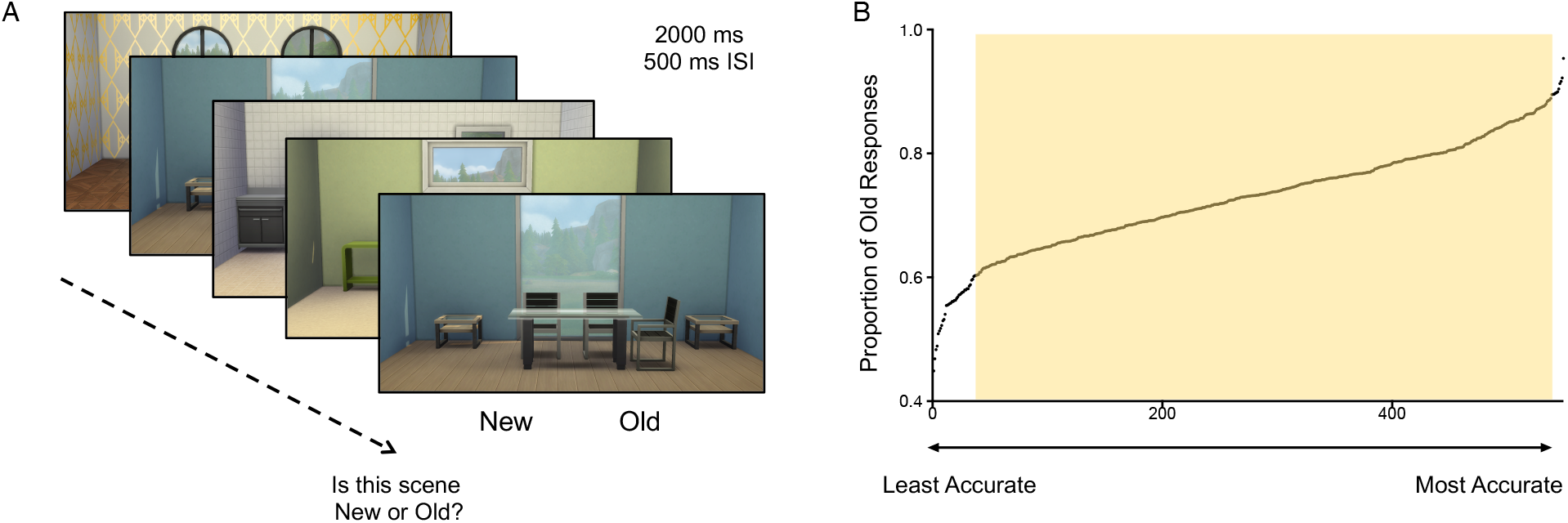
Scene Rating Experiment. A) Participants were presented with the scenes and asked to judge if each scene was “New,” (seen for the first time) or “Old,” (a repeated scene). B) Proportion of correct old responses to repeated scenes for each scene ranked from least to most often accurately recalled. Highlighted portion (60% - 90% accuracy) of scenes was utilized in the subsequent experiment.

Repeated stimuli were spaced apart a minimum of 15 trials and maximum of 40 trials between first, second and third presentations. Scenes were presented for 2500 milliseconds with a 500 milliseconds inter-stimulus interval. Stimuli were presented using Psychtoolbox 3 (Brainard, 1997; Pelli, 1997; Kleiner et al., 2007) using MATLAB (The Mathworks, Natick, MA) on a Macintosh computer.

## Results

Participants in experiment 1 correctly identified 67.9% of new trials as ‘new’, 76.1% of trials repeated once as old, and 85.7% of trials presented twice as ‘old’. To compare the accuracy between scenes, the average accuracy for each scene was computed by collapsing the correct trial type responses across participants for each scene. This resulted in an average accuracy for each scene for trials presented once, trials repeated once, and trials repeated twice (Figure 2B).

Of particular interest were the scenes judged ‘old,’ as this would be most representative of how reliably identifiable and memorable each of the scenes were. Average accuracy for trials repeated once and correctly called ‘old’ was used to compare the memorability of individual scenes. Scenes with accuracy scores between 60-90% were selected for use in subsequent experiments, resulting in a total of 509 scenes.

### Experiment 2: Object-in-Context

#### Participants

Twenty-eight young adults and nineteen cognitively normal older adults participated in the Object-in-Context experiment (Table 1). Young adults were recruited from the undergraduate population at Johns Hopkins University and received course credit for their participation. Older adults were recruited from the community through flyers and online advertisements and were paid for their participation. Data from twelve older adult and six young adult participants were excluded from analysis due to below chance performance (n = 15) or incomplete data collection (n = 3).

All older adult participants underwent medical, psychiatric, neurological and neuropsychological evaluations and completed the Clinical Dementia Rating Scale (CDR; Morris, 1993). Neuropsychological evaluation included the Mini Mental Status Exam (Folstein et al., 1975), the Buschke Selective Reminding Test (Buschke & Fuld, 1974), the Logical Memory subtest of the Wechslers Memory Scale (Wechsler, 1987), the Clock Drawing test (Sunderland et al., 1989), the Rey-Osterrieth complex figure test (Rey, 1941; Osterrieth, 1944), and the Benton Visual Retention Test (Benton, 1974). Participants were excluded from further participation if they reported current neurological or psychiatric disorders, history of major head trauma, history of substance abuse or dependencies, or scored 2.5 standard deviations below published norms on one or more neuropsychological tests. All older adult participants had a global CDR and CDR-Sum of Boxes score of 0.

#### Materials and Procedures

Stimuli consisted of 160 scene stimuli and 200 images of everyday objects. These scenes were randomly selected from the scenes validated in Experiment 1. Scenes were divided into 6 general categories: living room, dining room, kitchen, bedroom, or office rooms to allow for the placement of categorically relevant objects. All objects and scenes were categorically matched to allow coherency in the stimuli presentation (e.g., pencils in the office, apple in the kitchen).

#### Object Familiarization Phase

In the Object Familiarization phase of the experiment, participants viewed the 200 images of everyday objects and were asked to rate the objects as either belonging “Indoor” or “Outdoor” to familiarize themselves with all objects (Figure 3A). Each object was presented on a blank screen for 2.5 seconds with an inter-stimulus interval of 0.5 seconds.

**Figure 3.**
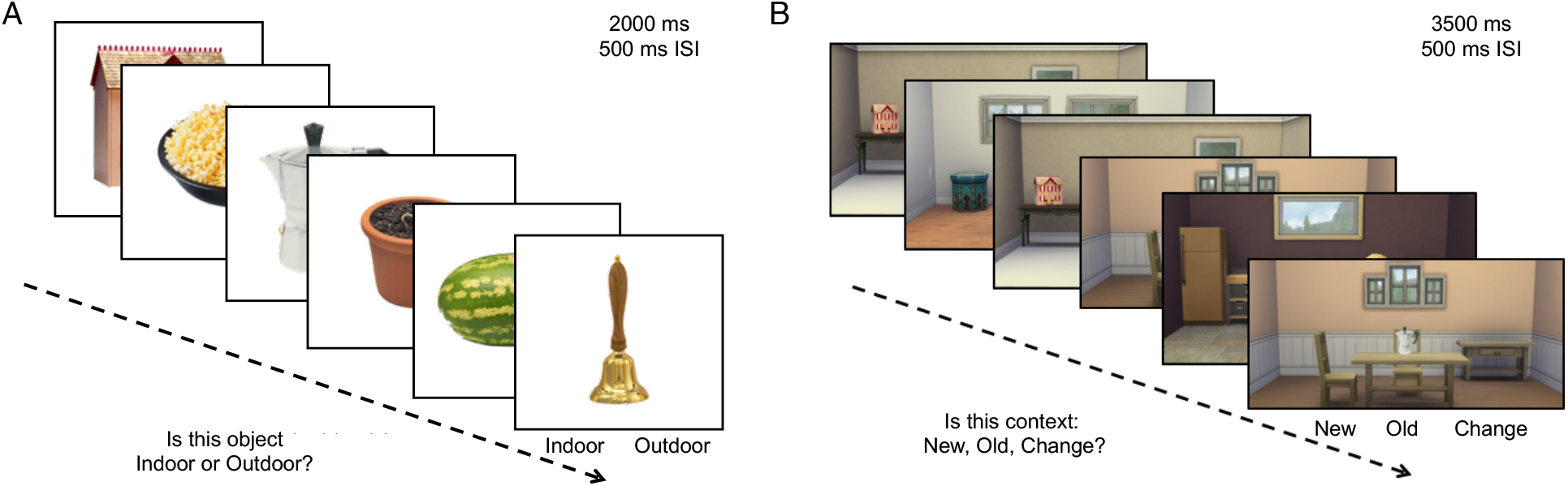
Object-in-Context Task. A) Participants were first presented with pictures of objects and asked to judge for each item if the object belongs “indoor” or “outdoor” to become familiar with the objects. B) Participants were subsequently presented with the scene and object pairing and asked to judge for each if the scene was “New” (never seen before), “Old” (previously seen) or “Change” (resembled a previously shown scene and object pair).

#### Object-in-Context Phase

In the Object-in-Context phase of the experiment, participants completed 280 trials where an object from the Object Familiarization phase was presented within a scene (Figure 3B). For each object-in-context trial, a scene was presented with a categorically appropriate object (e.g., blankets in a bedroom scene) for 3.5 seconds with an inter-stimulus interval of 0.5 seconds. Trials consisted of 40 “New” trials with new object and scene pairings that were presented once; 80 “Repeat” trials with 40 object and scene pairings that were presented and subsequently repeated once; 80 “Identity Change” trials with 40 object and scene pairings that were presented once and the object was changed on the subsequent trial using the same scene; and 80 “Position Change” trials with 40 object and scene pairings that were presented once and the position of the object was changed in a subsequent trial using the same scene. Participants were asked to rate a scene and object pairing as “New” when they were not seen before in the context of the task, “Old”, if the pairing was previously seen in the context of the task, or “Change”, for a previously seen pairing where either the object or the location of the object was changed. Repeated, Identity Change and Position Change trials were spaced apart a minimum of 3 trials and a maximum of 12 trials with an average of 8 trials between the first and second presentation.

Trials were presented in 6 different runs consisting of 70 trials per run. Runs were blocked so the first 3 runs would contain “New,” “Repeat”, and “Identity Change” trials while the last 3 runs would contain “New”, “Repeat”, and “Position Change” trials. This order was counterbalanced between participants. Participants were cued to the trial type at the beginning of each run. Previous versions of the task using no cues or a random order of trial types resulted in chance performance in older adults.

Stimuli were presented using Psychtoolbox 3 (Brainard, 1997; Pelli, 1997; Kleiner et al., 2007) using MATLAB (The Mathworks, Natick, MA) on a Macintosh computer.

## Results

Demographic data and neuropsychological test performance is included in Table 1. Older adults were by criteria significantly older than young adults (t(45)=47.54, p < 0.0001) and completed significantly more years of education (t(45)=6.84, p < 0.001). During the experimental task, young adults correctly identified 82.71% (SD = 13.07%) of New Trials while older adults correctly identified 62.83% (SD = 12.37%) of New trials (new object and scene pairing). For Repeat trials, where the same object and scene pairing are repeated, young adults correctly identified 82.40% (SD = 10.65%) of trials while older adults correctly identified 76.46% (SD = 15. 34%) of all trials. For Identity Change trials, with a new object and scene pairing, young adults correctly identified 66.56% (SD =16.55%) of trials while older adults correctly identified 74.24% (SD = 16.07%) of trials. For Position Change trials, where the position of the object is changed, young adults correctly identified 80.36% (SD = 13.11%) of trials while older adults identified 65.02% of trials (SD = 13.04%)

Older adults showed significantly poorer performance identifying new object-in-context trials (New trials: t(45) = 5.23, p < 0.001) and trials in which the position of the object in the context was changed (Position Change trials: t(45) = 3.95, p < 0.001) when compared to young adults. No significant differences were observed between young and older adults for trials in which the same object in context was repeated (Repeat trials: t(45) = 1.57, p = 0.12) or in which the object in the context was changed (Identity Change Trials: t(45) = 1.58, p = 0.12) (Figure 4).

**Figure 4.**
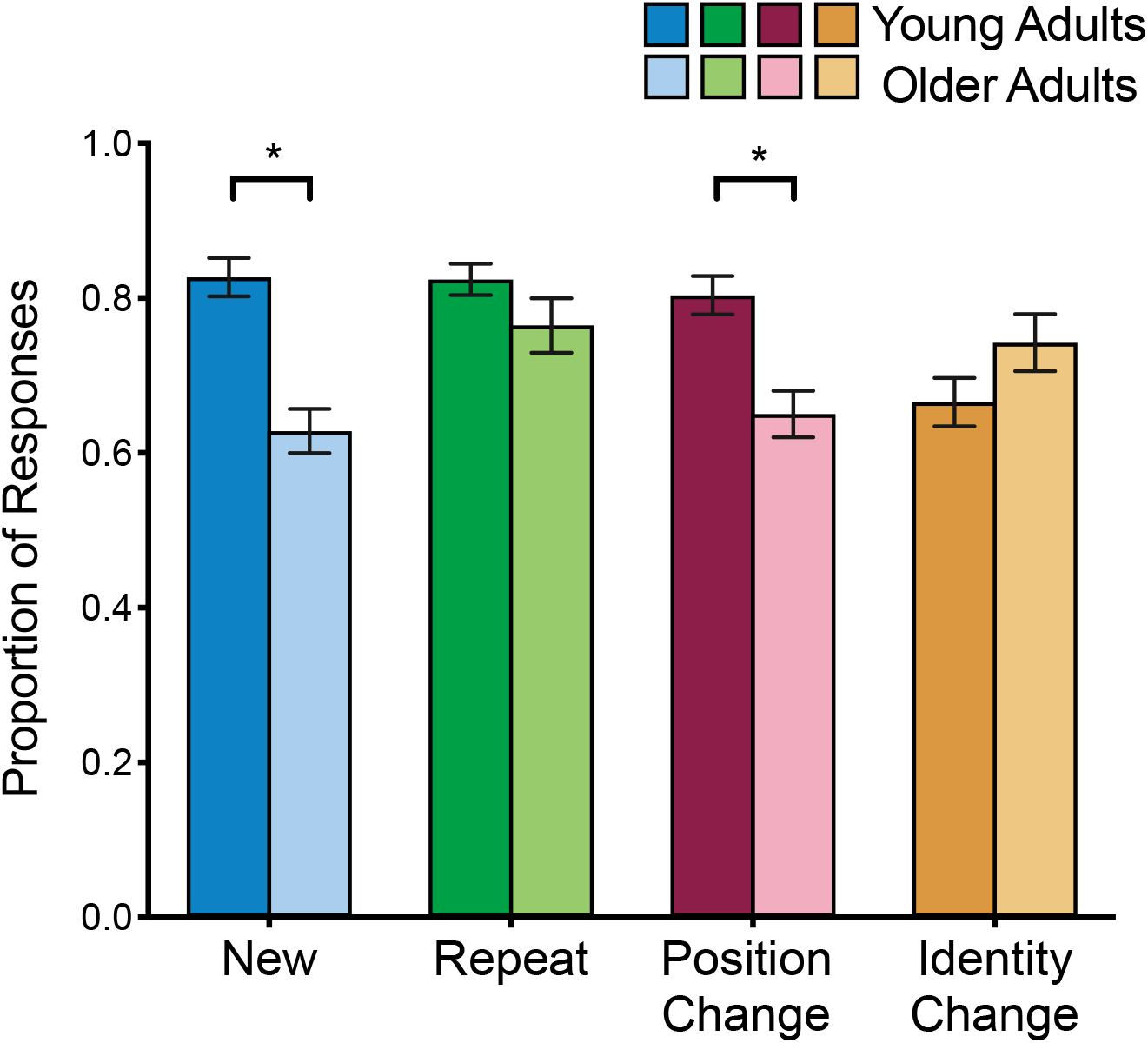
Behavioral performance in young and older adults. Older adults show impaired performance on Position Change trials and New trials relative to young adults. Bars represent mean +/−SEM. (* p < 0.05).

A key goal of the current study was to examine the potential dissociation and age-related changes in memory for object position changes compared to object changes when presented in the context of a scene. A two-way analysis of variance comparing the Identity Change and Position Change trials between young and older adults showed no main effect of age (F(1,45) = 0.63, p = 0.43) or condition (F(1,45) = 1.324, p = 0.26). However, the analysis showed a significant interaction between age and condition (F(1,45) = 15.86, p < 0.001), illustrating that older significantly less often correctly identified Position Change trials compared to young adults (Figure 5A). A subsequent analysis of variance was conducted to examine the response rates of each response option (new, old or change) for the Position Change and Identity Change conditions respectively. For the Identity Change trials, there was no significant interaction between young adults and old adults for the response type (F(1,45) = 0.49, p = 0.49) (Figure 5B). In contrast, for the Position Change trials, there was a significant interaction between young adults and older adults for the response type (F(1,45) = 19.20, p < 0.001), showing that older adults significantly more often incorrectly identified Position change trials as “Old” and less often correctly identified those trials as ‘New’ when compared to younger adults (Figure 5C).

**Figure 5.**
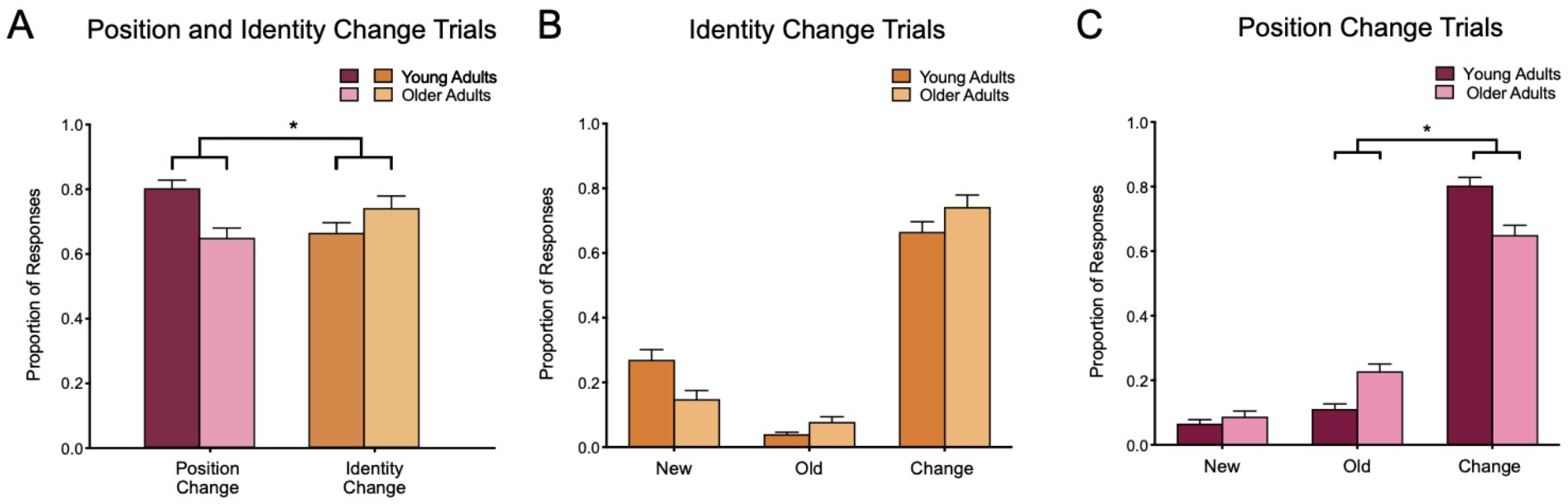
Older adults show impaired performance on Position Change but not Identity Change trials. A) Older adults showed a significant impairment on Position trials relative to young adults but no significant difference on Identity Change trials. B) Identity Change trials showed no significant difference in performance between older adults and young adults. C) For Position Change trials, older adults more often incorrectly identified position change trials as “old,” instead of “change” compared to young adults. Bars represent mean +/−SEM. (* indicate p < 0.05).

## Discussion

The goal of the current study was to assess age-related changes in the memory for objects and their position in a scene. Using a novel task for viewing different environments (living rooms, bedrooms, kitchens, dining rooms, and office rooms), younger and older adults were asked to identify a change in the identity of objects, as well as the position of the objects. Older adults correctly identified a similar number of object changes in the scenes but identified significantly fewer object position changes compared to young adults. These findings expand upon previous studies examining age-related differences reporting that older adults have difficulty recognizing changes in object identity (Olsen et al., 2017; Reagh et al., 2016; Stark et al., 2015, 2013; Yassa et al., 2011) and object features (Yeung et al., 2017) but not in recalling changes in the positions of an object when items were presented on a blank screen (Reagh et al., 2016; Reagh et al., 2018). Studies examining age-related differences in memory for spatial information in scenes have reported mixed results with some reporting minimal age-related impairments in scene recognition (Fidalgo et al., 2016; Stark & Stark, 2017) while others have reported no age-related differences (Berron et al., 2019, 2018). The majority of these studies examined object identity, scenes or an object’s position independently, without assessing the conjunctive encoding of an object and its position within a scene. However, mounting evidence suggests that object and position information, is encoded in relationship to the environmental context (Chao et al., 2016; Van Cauter et al., 2013; Wilson, Langston, et al., 2013).

As noted in the background for the current investigation, in foraging rodents, cells in the lateral entorhinal cortex track the position of an object in the environment (Deshmukh & Knierim, 2011) or the previous positions of an object within the environment (Tsao et al., 2013). Furthermore, a population-level analysis of neurons in the lateral entorhinal cortex shows that cells in this area encode for object, object position and context, and appear sensitive to encoding contextual information about the environment (Keene et al., 2016). These findings suggest that the representation of complex environments composed of the integration of spatial and non-spatial may be differentially processed relative to independent and isolated representations of objects and spatial information providing a potential explanation for the mixed results previously reported.

The task used in this study was designed to provide consistent spatial dimensions and perspective with objects congruent with the scene placed in plausible locations in the space. The scenes were equated for memorability based on ratings from a separate experiment in an effort to minimize the effects of novelty, perspective, congruence, and distance on recognition of object identity and object position. This task is similar to the approach employed in a study by Yeung and collogues showing that patients with cognitive decline are impaired in recognizing position changes in a scene relative to cognitively normal older adults (Yeung et al., 2019). In Yeung et al. (2019) the authors manipulated the position of an object in the environment, but with much larger changes including the presence of multiple new objects within the scene or moved elements within a scene. Performance on object position memory was correlated with anterolateral cortex and perirhinal cortex volume.

In contrast, the task utilized by Berron and colleagues (2018) and Reagh and colleagues (2018) featured an object mnemonic discrimination task, requiring discrimination between highly similar and overlapping representations of object identity or objection position on a blank screen without integration of spatial and non-spatial information. Despite the differences in the tasks employed, these studies have observed selective engagement of the entorhinal cortex consistent with the findings from recording studies in animals. Changes to object identity have been associated with activation of the lateral entorhinal cortex (Berron et al., 2018; Reagh et al., 2016, 2018) while changes to the position of the object (Reagh et al., 2016, 2018) and context (Berron et al., 2018) have been shown to elicit activation in the medial entorhinal cortex. Additional studies using neuroimaging approaches in humans are needed to determine if mechanisms underlying the representations of complex environments and whether the integration of spatial and non-spatial information also engage the lateral or medial entorhinal cortex or may be differentially processed relative to independent and isolated representations of objects and spatial information.

In addition to identifying significantly fewer position change trials, older adults also identified significantly fewer new scene and object pairings compared to young adults. This finding is consistent with reports of increased false recognition in older adults showing that older adults are more likely to falsely recall seeing words and pictures (Dennis & Turney, 2018; Koutstaal & Schacter, 1997; Norman & Schacter, 1997) than younger adults (Devitt & Schacter, 2016). Although increased false recognition in older adults has been well established, the mechanisms behind this phenomenon are not well understood and attributed to a general decline in prefrontal cortex integrity and connectivity between the prefrontal cortex and the medial temporal lobe (Devitt & Schacter, 2016).

Within the medial temporal lobe (MTL), there has been an increased focus on the lateral entorhinal cortex in recent years due to the vulnerability of this region in both aging and Alzheimer’s disease (AD). The lateral entorhinal cortex is one of the first regions where tau neurofibrillary tangles, an established classic biomarker of AD, accumulates (Braak & Braak, 1991; Jack et al., 2010; Lace et al., 2009), and subsequent neuronal degeneration and synaptic loss occurs (Gómez-Isla et al., 1996; Hoesen et al., 1991; Kordower et al., 2001; Selkoe, 2002). In cognitively normal older adults, the lateral entorhinal cortex also shows considerable neurofibrillary tangle deposits compared to young adults (Braak and Braak, 1991), suggesting that even in healthy aging, the structure and function of this region may be altered. Indeed, reduced volume of the lateral entorhinal cortex has been observed in lower performing older adults at high risk for developing AD (Olsen et al., 2017; Yeung et al., 2017). Furthermore, hypoactivity of the lateral entorhinal cortex associated with poor object recognition has been observed in both older adults (Berron et al., 2018; Reagh et al., 2018) and in patients with amnestic mild cognitive impairment, a transitional stage between healthy aging and AD dementia (Tran et al., under review). Using novel stimuli consisting of objects positioned within a scene, the current study shows there may be potential age-related differences in the mechanisms underlying the representations of complex environments. Furthermore, the integration of spatial and non-spatial information may be differentially processed relative to independent and isolated representations of objects and spatial information. It is possible that the integration of spatial and non-spatial information depends on the lateral entorhinal cortex and age-related changes are driven by the accumulation of pathology in this region in aging and AD. Further studies examining age-related changes and domain-specific processing are needed to determine the neural basis of information processing of these components in the medial temporal lobe.

## Acknowledgements

T.T. is supported by a NIA T32 training grant and a National Defense Science and Engineering Graduate Fellowship (NDSEG) grant awarded by the DoD, Air Force Office of Scientific Research, National Defense Science and Engineering Graduate (NDSEG) Fellowship, 32 CFR 168a. M.G. is the founder of AgeneBio. M.G. and A.B. are inventors on Johns Hopkins University intellectual property with patents pending and licensed to AgeneBio. M.G. consults for the company and owns company stock, which is subject to certain restrictions under University policy. M.G. and A.B’s role in the current study was in compliance with the conflict of interest policies of the Johns Hopkins School of Medicine.

